# Transcriptomic meta-analysis in plaque psoriasis: an integrative bioinformatic approach to deciphering the genetic landscape and molecular pathways

**DOI:** 10.1101/2025.03.01.640943

**Authors:** Teresa Torres-Moral, Josep Riera-Monroig, Gemma Tell-Martí, Jaume Bagué, José F. Català-Senent, Francisco J. Roig, Míriam Potrony, Francisco García-García, Susana Puig

**Affiliations:** Center for Biomedical Network Research on Rare Diseases (CIBERER), Carlos III Health Institute, Barcelona, Spain; August Pi i Sunyer Biomedical Research Institute (IDIBAPS), Barcelona, Spain; Dermatology Department, Hospital Clínic de Barcelona, University of Barcelona, Barcelona, Spain; Computational Biomedicine Laboratory, Principe Felipe Research Center (CIPF), Valencia, Spain; Faculty of Health Sciences, San Jorge University, Villanueva de Gállego Campus, Zaragoza, Spain; Biochemistry and Molecular Genetics Service, Hospital Clínic de Barcelona, Barcelona, Spain

**Keywords:** Plaque psoriasis, meta-analysis, transcriptomics, functional profiling, biomarkers

## Abstract

Psoriasis is a chronic inflammatory skin disease influenced by both genetic and environmental factors. Despite extensive research, its precise etiology remains unclear, posing significant challenges to understanding and treatment. The disease pathogenesis involves self-reactive T cells and immune-related cytokines. Genome-wide association studies have identified various susceptibility loci for immune-related diseases, but the underlying mechanisms remain only partially understood. Recent discoveries of critical signaling pathways, biological processes, and immune cell involvement have expanded our knowledge and offer hope for improved therapeutic strategies.

This study aimed to enhance our understanding of psoriasis and proposes novel therapeutic approaches by employing integrated bioinformatics to identify signaling pathways and biological processes as potential disease markers.

Presenting a systematic review and taking a meta-analytical approach to transcriptomic profiles, this investigation examined differential gene expression patterns across 44 studies involving 975 samples comparing lesional psoriasis, non-lesional psoriasis, and healthy controls. Consensus transcriptome signatures revealed a significant association between immune-related genes and psoriasis pathogenesis. Functional enrichment analysis identified several enriched pathways related to immunity and immune system processes. Comparison of these findings with the existing literature indicated that some immune-related genes were already known, while others are novel in the context of psoriasis. Additionally, novel gene analysis demonstrated psoriasis involvement in pathways such as *gluconeogenesis*, the *FoxO signaling pathway*, and *mitophagy*.

This integrative approach confirmed classic genetic associations while uncovering novel gene expression patterns and pathways relevant to psoriasis. Notably, the disruption of the *gluconeogenesis* pathway emerged as a critical finding. These insights enhance our understanding of psoriasis pathophysiology and pave the way for targeted therapies, offering improved management options for affected individuals.

## Introduction

Psoriasis is a chronic, immune-mediated inflammatory skin disease that manifests in a range of severities, from scattered red, scaly plaques to widespread involvement across the body. This condition is known to be influenced by various genetic and environmental factors, causing its severity to fluctuate over time. As a consequence, patients often experience a considerable impact on their quality of life, leading to psychosocial disability (Armstrong & Read, 2020; Parisi et al., 2013). Moreover, psoriasis is not solely confined to skin-related complications. It has been associated with several comorbidities including cardiovascular disease, fatty liver disease, and depression, as well as psoriatic arthritis: a multifaceted and complex condition difficult to manage effectively.

Despite advances in learning, the exact etiology of psoriasis remains elusive. However, several risk factors have been identified, shedding some light on potential causality. Family history plays a significant role, indicating a genetic predisposition to the disease. Furthermore, lifestyle factors such as smoking, stress, obesity, and alcohol consumption have all been linked to the onset and exacerbation of psoriasis symptoms (Armstrong & Read, 2020; Parisi et al., 2013).

Geographically, psoriasis prevalence varies, affecting approximately 2%–3% of the population in Western countries. However, this number may vary based on factors such as age, gender, ethnicity, and geographic location, and is likely influenced by genetic and environmental components. Additionally, discrepancies in study design and the criteria used for diagnosis may also contribute to variations in prevalence estimates (Armstrong & Read, 2020; Parisi et al., 2013). In summary, psoriasis represents a complex and diverse dermatological condition, with a significant impact on patients’ lives. Understanding its underlying causes and risk factors remains an ongoing challenge for researchers. Improved insights into the mechanisms of the disease will pave the way for better management and targeted therapies to alleviate its burden on affected individuals.

Although psoriasis is a complex inflammatory skin disease with multiple contributing factors, it is clear that the immune system plays a significant role (Wang et al., 2020). Specifically, autoreactive T cells and a network of cytokines are strongly implicated. It is well known that the IL-23/Th17 axis acts as a central signaling pathway providing crucial insights into the accelerated inflammation in psoriasis (Srivastava et al., 2021).

Regarding genetic background, numerous studies have focused on identifying psoriasis susceptibility loci that are associated with its pathogenesis. Large-scale case-control Genome-Wide Association Studies (GWAS) for psoriasis have revealed that many single-nucleotide polymorphisms (SNPs) linked to the condition are located within genes involved in adaptive and innate immune responses crucial for T helper 17 cell activation (Dand et al., 2023). These genes include those encoding the human leukocyte antigen (HLA)-C, interleukin IL-23 receptor, IL-23A, IL-12B, and TRAF3-interacting protein 2 (Ogawa & Okada, 2020; Wang et al., 2020; J. J. Wu et al., 2022). The relationship between the immune system and psoriasis is so significant that a recent study has proposed a bioinformatic gene signature score based on the mRNA of the immune landscape of psoriasis lesions. This score holds promise as a potential predictor of psoriasis treatment outcomes (Wang et al., 2020). In conclusion, the relationship between the immune system and psoriasis is strong, but also intricate and multifaceted.

The discovery of new critical signaling pathways, biological processes, and immune cell involvement has advanced our understanding of psoriasis pathogenesis and offers hope for improved treatment options and outcomes for affected individuals. In this article we present a novel integrative strategy based on a systematic review and meta-analysis of transcriptomic profiles in psoriasis which has allowed us to identify potential disease markers that improve our understanding of psoriasis, as well as to make proposals for new therapeutic approaches.

## Results

### Systematic review and selection of studies

Our systematic review screened 306 publications, which rendered a total of 44 studies eligible for our analyses (**Figure 1**). These selected studies contained 975 samples - 513 psoriasis patients and 462 healthy control patients (**Supplementary Table 1**).

**Figure 1.**
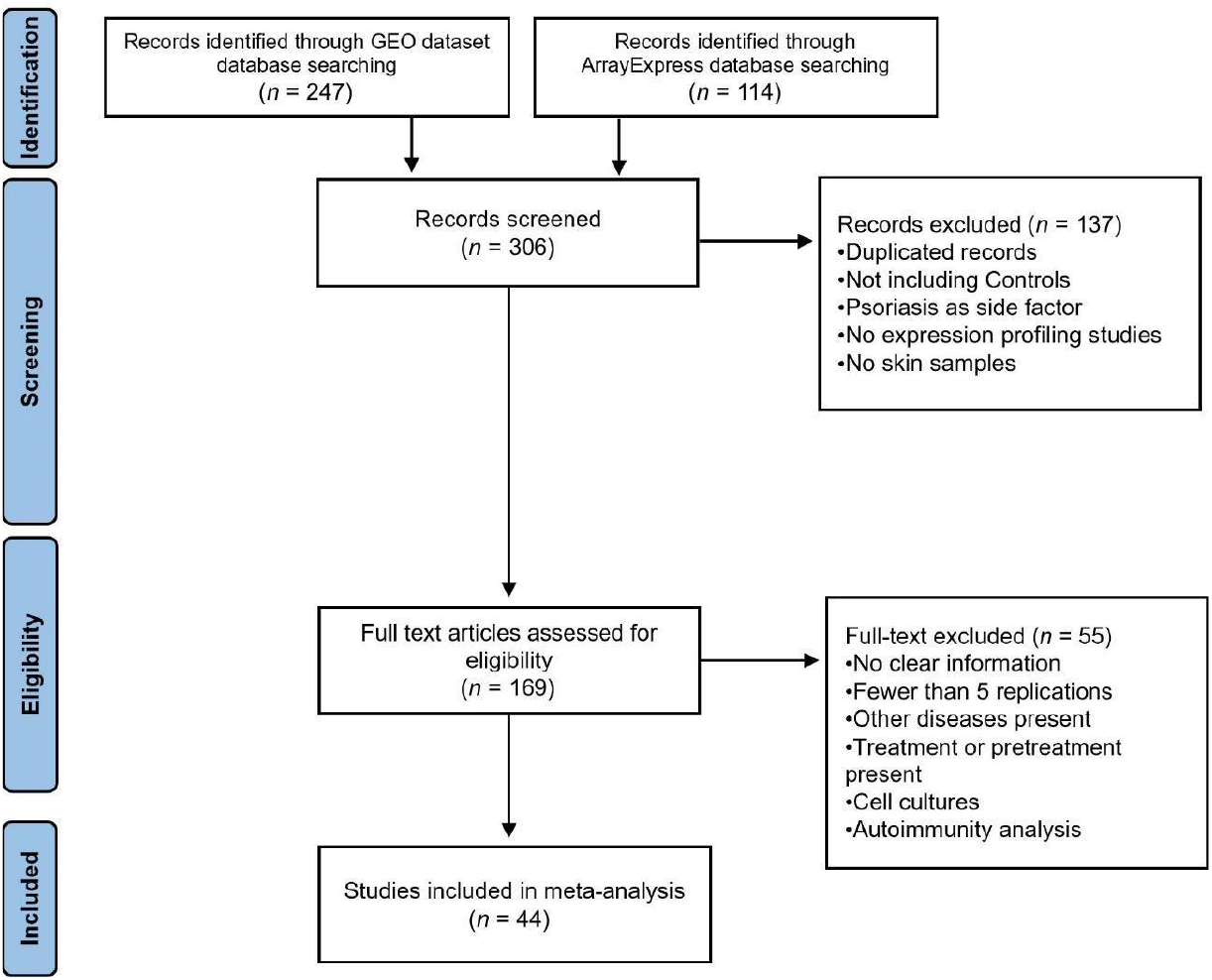
The flow of information through the distinct phases of the systematic review following PRISMA statement guidelines.

### Consensus psoriasis transcriptome signatures

For each of the selected studies, the data were downloaded, processed and normalized. Differential expression was then assessed for the comparisons included in each of these studies: a) lesional psoriasis vs. non-lesional psoriasis, b) lesional psoriasis vs. controls, and c) lesional psoriasis vs. non-lesional psoriasis plus controls.

Subsequently, the set of expression results from the selected studies was integrated using a transcriptomic meta-analysis strategy to identify consensus expression signatures in the comparisons evaluated. From the comparison of lesional psoriasis vs non-lesional skin from the same patient, we identified data expression from 2,385 genes related to the immune system, of which 1,673 were found to be significant. When comparing lesional psoriasis with controls, we found data expression from 2,534 immune system genes, of which 1,328 were significant. In the comparison of lesional psoriasis with non-lesional psoriasis plus controls, data expression from 2,538 immune system genes was identified, where 1,780 were significant (**Figure 2, Figure 3**). To ensure the inclusion of all immune system genes relevant to psoriasis, we selected the 1,780 significant genes identified in the comparison of lesional psoriasis with non-lesional psoriasis plus controls for later analysis (**Supplementary Table 2**).

**Figure 2.**
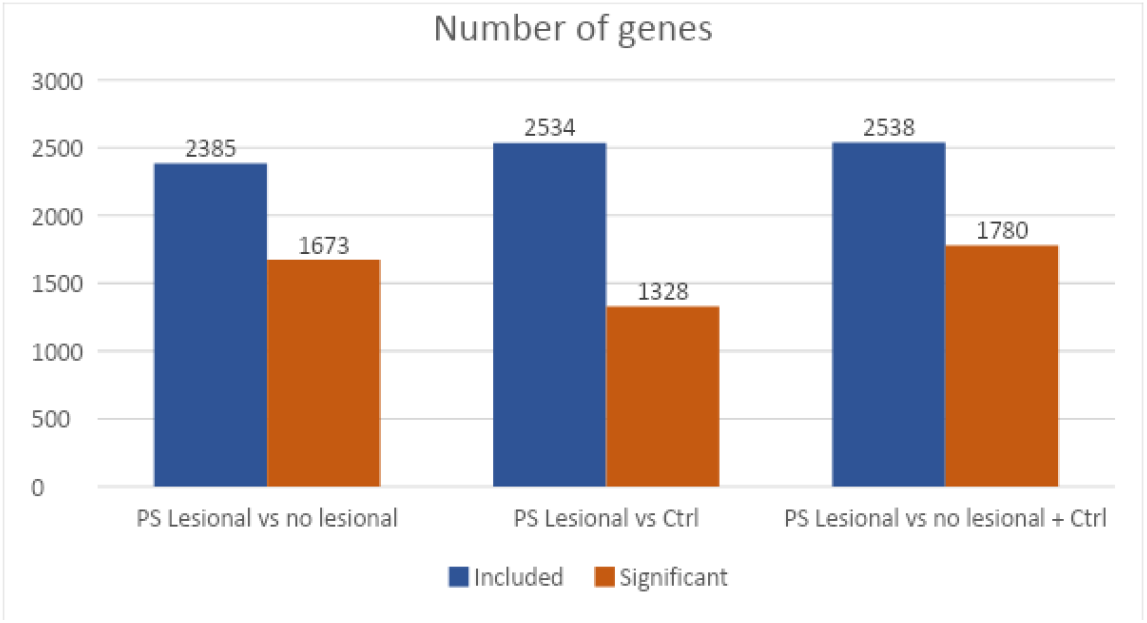
Number of included and statistically significant genes for each comparison in our meta-analysis: lesional psoriasis vs. non-lesional psoriasis; lesional psoriasis vs. controls, and lesional psoriasis vs. non-lesional psoriasis plus controls.

**Figure 3.**
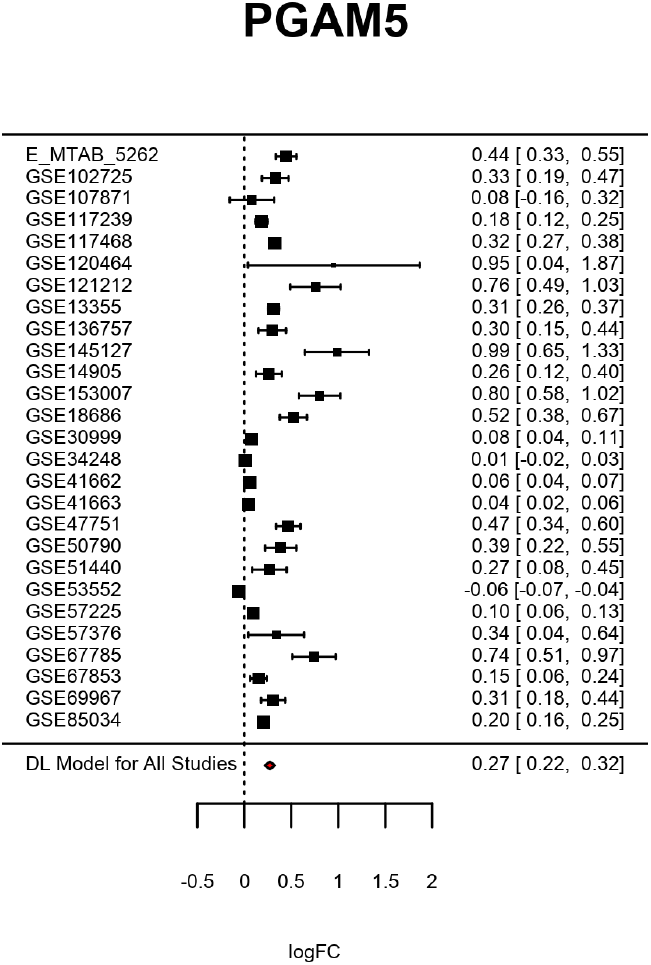
Meta-analysis results. Representative forest plot showing the result of the *PGAM5* gene result for the meta-analysis of lesional psoriasis vs. nonlesional psoriasis samples. On the left, for each study, the change in expression between the two groups compared for the *PGAM5* gene is represented by the logarithm of the fold-change (logFC) and its confidence interval. At the bottom of the graph, the logFC obtained after applying the random-effects meta-analysis that summarises the individual results into a single consensus measure is shown.

### Functional enrichment analysis

A functional enrichment analysis was performed on the 1,780 significant genes identified in the comparison of lesional psoriasis with non-lesional psoriasis plus controls, based on three ontologies: Reactome, KEGG, and Biological Process Gene Ontology (BP-GO). We selected those results with a BY adjusted p-value < 0.05 and including at least 10 genes, obtaining 31 pathways in the Reactome database, 11 in KEGG, and 203 in BP-GO. The detailed results are presented in **Figure 4** and **Supplementary Table 3**.

**Figure 4.**
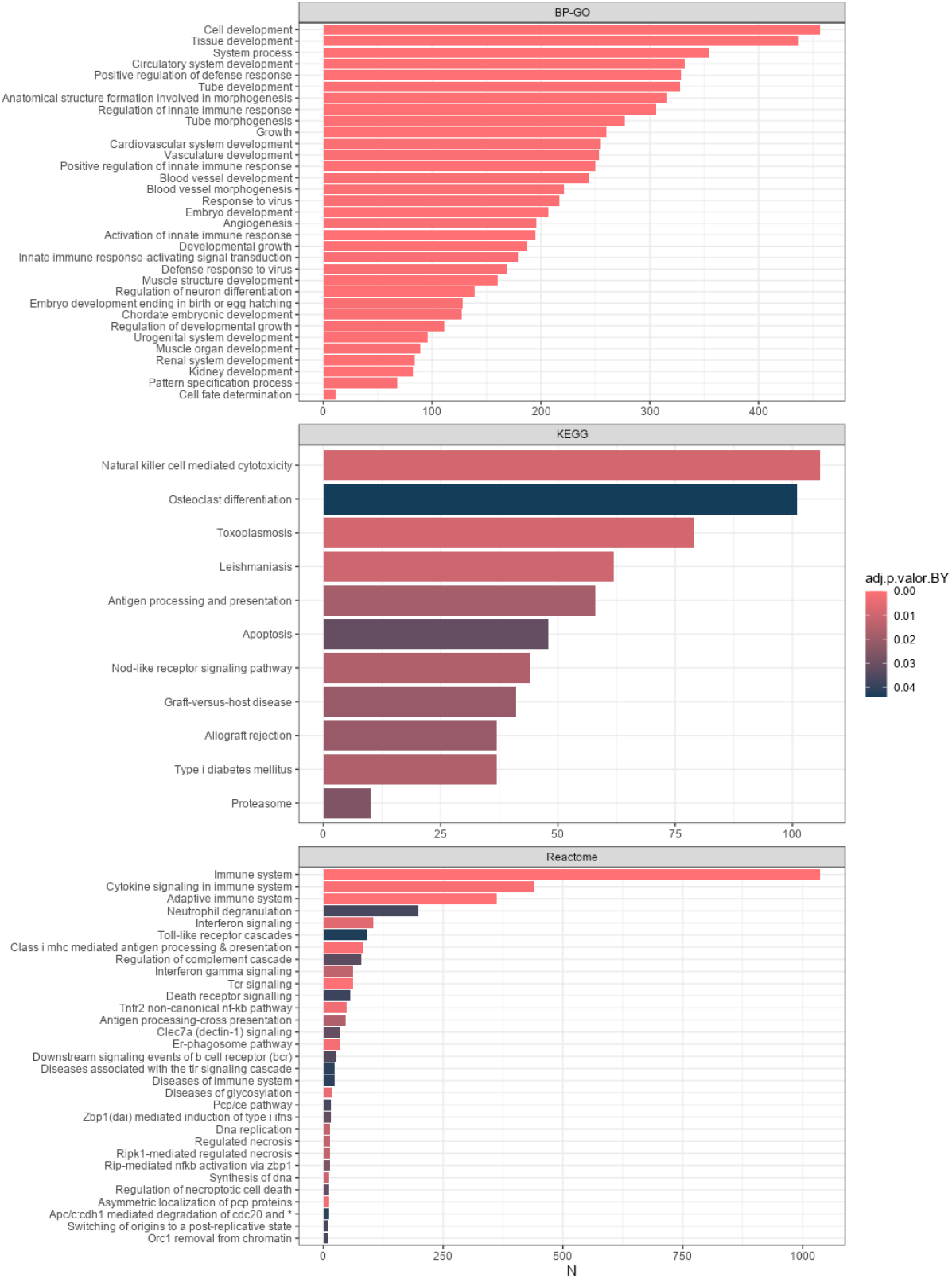
Significant results from functional enrichment analysis. **A)** Biological processes of Gene Ontology; **B)** KEGG pathways; **C)** Reactome pathways.

Our search in the Open Targets Platform (OT) yielded 6,196 genes related to psoriasis, with 1394 of them related to the immune system. When comparing this with our 1,780 immune DEGs from our meta-analysis, we found a coincidence of 1,119 genes, accounting for 62.86% of the DEGs from our meta-analysis, and 661 genes specific to our meta-analysis (**Supplementary Table 2**).

### Functional profiling in newly associated genes

We identified 157 (8.8% of 1,780) significant genes from our meta-analysis that were not associated with any of the previously identified enriched pathways, nor have they been previously reported in OT. We conducted a gene enrichment analysis of these genes using the same databases as in the previous functional enrichment analysis (Reactome, KEGG, and biological processes from Gene Ontology), resulting in a total of 5 significant pathways: *gluconeogenesis* from BP-GO, *FoxO signaling* pathway from KEGG and *Mitophagy, PINK1-PRKN mediated mitophagy* and *Selective autophagy* from Reactome (**Table 1**).

**Table 1.** Functional profiling in new associated genes. Significant pathways found from our meta-analysis in significant genes, which were not associated with any of our previously identified enriched pathways or previously reported in Open Targets Platform.

| Database | Pathway name | Name | Adjusted p-value | Genes included |
| --- | --- | --- | --- | --- |
| BP-GO | GO:0006094 | *Gluconeogenesis* | 0.039352 | 3 (*G6PC3*, *TPI1*, *USP7*) |
| KEGG | KEGG:04068 | *FoxO signaling pathway* | 0.048127 | 5 (*AGAP2*, *CCND2*, *G6PC3*, *GADD45G*, *USP7*) |
| Reactome | R-HSA:5205647 | *Mitophagy* | 0.001184 | 4 (*MAP1LC3B*, *PGAM5*, *TOMM40*, *TOMM70*) |
| Reactome | R-HSA:5205685 | *PINK1-PRKN mediated mitophagy* | 0.009582 | 3 (*MAP1LC3B*, *TOMM40*, *TOMM70*) |
| Reactome | R-HSA:9663891 | *Selective autophagy* | 0.044155 | 4 (*MAP1LC3B*, *PGAM5*, *TOMM40*, *TOMM70*) |

## Material and methods

### Selection of gene expression datasets

This systematic review of the Gene Expression Omnibus (GEO) and ArrayExpress databases (Barret et al., 2013; Sarkans et al., 2018) was performed with the keywords SKIN, PSORIASIS, DERMIS and EPIDERMIS. The period reviewed ranged from January 2002 (generation of the public repositories) to May 2020. Results were filtered by: (1) organism: ‘Human’, (2) study type: ‘expression profiling by array’ or ‘expression profiling by high throughput sequencing’. To ensure the quality and transparency of the systematic review, PRISMA statement guidelines were followed (Page et al., 2021). In addition, the following exclusion criteria were applied: (1) information about inclusion criteria not available, (2) transcriptomic studies in which only non-coding RNA was analyzed.

#### Data acquisition and preprocessing

As all data used for this study were publicly available and completely deidentified, the study was exempt from any institutional review board approval by Hospital Clinic of Barcelona. We conducted a systematic review benchmarking published psoriasis gene expression data sets available on the Gene Expression Omnibus (https://www.ncbi.nlm.nih.gov/geo/) and selected data sets with 5 or more samples per experimental group. The database searches were conducted in January 2018 and updated in April 2020.

Expression data from selected studies in the systematic review were downloaded and standardized. The gene names were in accordance with Gene Symbol nomenclature and t he median expression was calculated when multiple values were obtained for the same gene. Individuals were categorized into three groups: lesional psoriasis, non-lesional psoriasis and controls. After normalization of the data, an exploratory analysis was performed to detect possible batch effects and anomalous behavior within the data, by means of clustering and principal component analysis.

#### Differential Gene Expression (DGE) and Meta-analysis

After normalizing the data within each study, we conducted the differential expression gene analysis, comparing lesional psoriasis vs. non-lesional psoriasis, lesional psoriasis vs. controls, and lesional psoriasis vs. non-lesional psoriasis plus controls. The methods used were limma (Ritchie et al., 2015) and *DEseq2* (Love et al., 2014) for microarray and RNA-seq studies, respectively. Additionally, a selection of 2,583 genes related to the immune system was conducted. To identify immune-related genes, we used a two-step approach. First, we curated the list of differentially expressed genes by selecting those associated with Gene Ontology (GO) (Gene Ontology Consortium et al., 2023) terms related to the immune system. Next, to capture genes potentially involved in the immune response but lacking assigned GO annotations, we further cross-referenced individual gene descriptions with terms such as “T cells”, “lymphocytes”, “Helper T cells”, “Th cells”, “Cytokines”, “Innate immunity”, and “autoimmunity”. P-values were adjusted by the Benjamini & Hochberg method (Benjamini and Hochberg, 1995). Genes with an adjusted p-value < 0.05 were considered significantly differentially expressed (DEGs). All analyses were conducted using R version 3.6.2 (R Core Team, 2023).

To robustly integrate the DGE results of each study, a meta-analysis technique was applied to each one of the comparisons defined above. The meta-analyses were performed following the methodology described in García-García (2016). Briefly, the *metafor* package for R (Viechtbauer, 2010) was used to combine the statistics from each of the individual studies, using a random effects model for each gene, which is particularly suitable when there are several sources of variability (DerSimonian and Laird, 1986). The meta-analyses calculate a pooled measure of the expression effect across all studies for each gene (pooled logFC and its 95% confidence interval) and provide a BH-adjusted p-value (Benjamini and Hochberg, 1995), with a significance threshold set at 0.05.

With the available gene expression data, our first step was to collect all genes with information related to the immune system. Subsequently, we conducted the differential expression gene analysis, comparing lesional psoriasis with non-lesional psoriasis, lesional psoriasis with controls, and lesional psoriasis with non-lesional psoriasis plus controls. Genes with an adjusted p-value < 0.05 were considered significantly differentially expressed (DEGs). All analyses were conducted using R version 3.6.2.

### Functional enrichment analysis

Based on the results of the meta-analysis, we performed a functional enrichment analysis using the *mdgsa* R package (Montaner & Dopazo, 2010). Three ontologies were analyzed: Reactome (Jassal et al., 2020, https://reactome.org), KEGG (Kanehisa et al., 2000, https://www.genome.jp/kegg/), and the biological processes category of Gene Ontology (BP-GO) (Ashburner et al., 2000, http://geneontology.org/). Pathways or biological processes with a Benjamini & Yekutieli (2001) (BY) adjusted p-value < 0.05 and containing at least 10 genes were considered significant.

### Literature comparison and functional profiling in new associated genes

On September 17, 2021, a search for genes previously associated with psoriasis was performed using the Open Targets Platform (Carvalho-Silva et al., 2019). This search aimed to compare the immune system-related genes identified in our meta-analysis with those reported in the literature.

To identify novel pathways and biological processes involved in psoriasis, we focused on significant genes from our meta-analysis that were not registered in Open Targets (OT) and were not associated with pathways or processes identified as significant in our initial functional enrichment analysis. For these genes, we performed a functional enrichment analysis using the same databases as before: Reactome, KEGG, and the biological processes category of Gene Ontology. Pathways with a Bonferroni-adjusted p-value < 0.05 were considered significant and included in the analysis.

## Discussion

This study aimed to investigate the role of the immune system in the development of lesional psoriasis by comparing its gene expression with that of healthy skin. The primary focus was to gain a deeper understanding of novel pathways, biological processes and genes involved in the pathogenesis of psoriasis.

After conducting the meta-analysis, we identified a substantial number of datasets and genes associated with psoriasis lesions. These findings align with information gathered from previous research projects aimed at understanding the illness, motivated by its high prevalence of 2%–4% in Western countries. (Armstrong & Read, 2020; Parisi et al., 2013)

The analysis of all significant immune system genes revealed a considerable number of significant pathways (275). Among these pathways, the most significant were related to the Immune System and Adaptive Immune System. These findings are consistent with our selection criteria and align with previous literature on the subject. (Armstrong & Read, 2020; Parisi et al., 2013)

Analyzing the results, we discovered that more than 10% of the significant genes identified in our meta-analysis had not previously been reported as associated with psoriasis. These newly identified genes showed enrichment in five pathways, none of which overlapped with the pathways found in all the significant immune system genes. This underscores the importance of a substantial sample size and collaborative efforts in identifying relevant genes and pathways in psoriasis. The pathways identified through our analysis include *gluconeogenesis*, the *FoxO signaling pathway, mitophagy*, and *PINK1-PRKN mediated mitophagy* pathways, together with *selective autophagy*.

Gluconeogenesis is the process through which specific molecules, such as lactate, pyruvate, glycerol, and amino acids, are converted into glucose and glycogen. To the best of our knowledge, a direct relationship between gluconeogenesis and psoriasis has not been clearly defined. However, it is known that psoriasis is associated with various metabolic alterations, including obesity, insulin resistance, and diabetes. Since gluconeogenesis is a metabolic pathway, it may play a role in the utilization of amino acids, which could be released during protein breakdown in tissues affected by psoriasis (Christophers, 2001; Gupta & Gupta, 2021; Heidenreich et al., 2009). Based on the information presented, we propose that the genes involved in the gluconeogenesis pathways may be genetically altered in patients with psoriasis. This genetic alteration could potentially impact skin cells and lead to an increased protein breakdown in affected tissues. However, it is essential to note that further research is required to comprehensively understand the potential link between gluconeogenesis and psoriasis.

In the *gluconeogenesis* pathway, we identified *G6PC3, TPI1*, and *USP7. G6PC3* gene mutations have been linked to disrupted glucose metabolism and heightened endoplasmic reticulum stress, which, in turn, might contribute to neutrophil dysfunction (Hayee et al., 2011; Smith et al., 2012; Xue et al., 2022). We propose that this deficiency in the *gluconeogenesis* pathway may potentially play a role in the pathogenesis of psoriasis. As for *TPI1*, it encodes for triosephosphate isomerase, which catalyzes the conversion of dihydroxyacetone phosphate to glyceraldehyde 3-phosphate and has been related to cancer progression (Jin et al., 2022; Valentin et al., 2000). TPI deficiency is a severe syndrome characterized by hemolytic anemia, neurological abnormalities and increased risk of infection. Regarding psoriasis, TPI1 has been shown to be overexpressed in CD8+ T cells exhibiting high dysfunction in both psoriasis and melanoma (Liu et al., 2021). *USP7* is a deubiquitinating enzyme known to play a crucial role in regulating the stability and expression of various proteins by removing ubiquitin molecules. It has been shown to interact with and deubiquitinate proteins such as Acf7, NHE3, p57KIP2, and myogenin, among others (de la Vega et al., 2020; Han & Yun, 2020; Yan et al., 2021). Clearly, ubiquitination is a critical cellular process that modulates various biological activities, encompassing immune responses, protein degradation, apoptosis, T cell activation and differentiation, and protein localization. It also plays a crucial role in regulating central and peripheral immune tolerance to prevent the development of autoimmunity. Any deregulation in the ubiquitination proteasome system can lead to aberrations in inflammation-related pathways, potentially contributing to the occurrence of autoimmune diseases, including psoriasis. Furthermore, it is worth noting that mutations in several E3 ligases and deubiquitinases have been identified and characterized in psoriasis patients. (Yang et al., 2018)

The *FoxO pathway* mediates the forkhead box O (FOXO) transcription factors, and inhibiting FoxO reduces the levels of p27Kip1 and IκB, promoting cell cycle progression and inhibiting cell proliferation (Dorai & Alex Anand, 2022; Zhang & Zhang, 2019). Another recent study has shown that, in the non-lesional skin of psoriatic patients, a cell-cycle inhibitory process acts as a stress-related compensatory mechanism, blocking the pathway to prevent keratinocyte hyperproliferation (Bozó et al., 2021).

FoxO is negatively regulated by the *PI3K/AKT signaling pathway*, which plays a central role in multiple cellular functions, such as cell proliferation and survival. In psoriasis, characterized by hyperproliferative keratinocytes, dysregulation of the *FoxO pathway*’s genetic expression likely contributes to this keratinocyte hyperproliferation (Zhang & Zhang, 2019). Further investigating the role of the *FoxO/PI3K/AKT signaling pathway*, another study highlights the downregulation of miRNA-559 as a promising diagnostic biomarker, inducing proliferation and inhibiting apoptosis through the *PTEN/AKT pathway* via positive regulation of metadherin MTDH (Aldabbas et al., 2023).

Regarding mitophagy, specifically the *PINK1-PRKN mediated mitophagy* pathway, it is known that the pro-inflammatory cytokine TNF is a key driver of psoriasis, and TNF inhibits PINK1-mediated mitophagy, leading to altered mitochondrial function (Willemsen et al., 2021). Furthermore, recent studies have demonstrated that autophagy also regulates highly selective degradation processes, such as clearing damaged mitochondria through mitophagy.

Regarding autophagy, it is known to have a negative regulatory effect on TLR2/6 mediated NF-κB activation, *SQSTM1* expression, and cytokine secretion in human keratinocytes, which are critical factors contributing to skin inflammation observed in psoriasis (D. J. Wu & Adamopoulos, 2017). Chronic skin inflammation, especially long exposure to TNF-a and IL-17A, has been linked to a decrease in autophagy, and in turn, this alteration in autophagy can lead to aberrant keratinocyte differentiation and increase cytokine production (Klapan et al., 2022). Polymorphisms on autophagy gene *ATG16L1* have been linked to a higher risk of developing psoriasis (Douroudis et al., 2012). Because of the close relationship between dysfunctional autophagy processes and inflammatory skin disorders, some authors have postulated its potential role as a therapeutic target (Hailfinger & Schulze-Osthoff, 2021).

In summary, this study conducted a meta-analysis of the role of immune system genes in psoriasis. The analysis reinforces the genetic associations of certain genes with their related biological pathways. Additionally, it uncovers new genetic and pathway associations implicated in psoriasis, with particular significance given to the disruption of the *gluconeogenesis* pathway.

## Conflict of interest statement

The authors state no conflict of interest.

## Data availability statement

Transcriptomic datasets evaluated to perform meta-analyses are stored in the Gene Expression Omnibus or Array Express databases with accession numbers: E_MTAB_5262, GSE102725, GSE107871, GSE109248, GSE114286, GSE117239, GSE117468, GSE11903, GSE120464, GSE121212, GSE13355, GSE136757, GSE145127, GSE14905, GSE153007, GSE18686, GSE26952, GSE30999, GSE32407, GSE34248, GSE41662, GSE41663, GSE47751, GSE50790, GSE51440, GSE52471, GSE53431, GSE53552, GSE57225, GSE57376, GSE58121, GSE63741, GSE6710, GSE67785, GSE67853, GSE69967, GSE75343, GSE75890, GSE78097, GSE79704, GSE80047, GSE83582, GSE83645, GSE85034.

## Acknowledgements

The authors express their gratitude to the Principe Felipe Research Center (CIPF) for providing access to the cluster, co-funded by the European Regional Development Funds (FEDER) for the Valencian Community 2014-2020. We also thank the Esther Koplowitz Center, IDIBAPS, Barcelona, where part of the work was conducted. This research was partially financed by the CIBER de Enfermedades Raras of the Institute of Health Carlos III, Spain, co-funded by ISCIII-Subdirección General de Evaluación, AGAUR 2021_SGR_01492, and the CERCA Programme by Generalitat de Catalunya, Spain. Moreover, this research received support and partial funding from the Institute of Health Carlos III (project IMPaCT-Data, IMP/00019 and PI18/00419), co-funded by the ERDF, ‘A way to make Europe’, and PID2021-124430OA-I00, funded by MCIN/AEI/10.13039/501100011033, CIAICO/2023/149 funded by the Consellería de Educación, Cultura, Universidades y Empleo de la Generalitat Valenciana, and by the ‘ERDF A way of making Europe’.

## Author contributions statement (CRediT-compliant)

Conceptualization: SP, FGG; Data Curation: FJR, JFCS; Investigation: TTM, JRM, GTM, JB, JFCS, FJR, NCL, MP, FGG, SP; Formal Analysis and visualization: FJR, TTM, JBC, JFCS, FGG; Supervision: SP, FGG; Writing-Original Draft Preparation: TTM, JRM, GTM, JB, JFCS, FJR, NCL, MP, FGG, SP; all authors read, review and approved the final manuscript.

## Notes

### Competing Interest Statement

The authors have declared no competing interest.

